# Extracellular tumor-derived IL-33 promotes ST2-dependent tumor control

**DOI:** 10.64898/2026.11.24.768800

**Authors:** Anja Herbst, Ihssane Eladnani, Ioannis Kritikos, Silvio Eugster, Vivian Pham Vu, Kristyna Saxena, Coline Nydegger, Marie-Hélène Wasmer, Robert Gaultney, Philippe Krebs

**Affiliations:** Institute of Tissue Medicine and Pathology, University of Bern, Bern, Switzerland; Graduate School for Cellular and Biomedical Sciences, University of Bern, Bern Switzerland

## Abstract

Interleukin-33 (IL-33) is a pleiotropic cytokine with both nuclear and extracellular functions as well as context-dependent roles in tumor immunity. Although tumor-derived IL-33 can promote anti-tumor immune responses, the contribution of its subcellular localization and extracellular availability to tumor control remains poorly understood. Here, we generated MC38 mouse adenocarcinoma cells overexpressing distinct IL-33 isoforms to dissect the relative contribution of IL-33 nuclear retention versus extracellular activity in a model of heterotopic tumors. Full length IL-33 (*FL*), an alanine-mutated isoform (*Ala*) with impaired chromatin binding and a truncated isoform (*Trunc*) of IL-33 lacking the nuclear functional domains were stably expressed in MC38 cells. These different IL-33 isoforms exhibited distinct subcellular localization and release profiles but had comparable effects on cell proliferation *in vitro*. *In vivo*, IL-33 overexpression reduced tumor growth compared with control MC38 cells, with the most pronounced growth impairment observed in tumors expressing *Trunc*. *Trunc* tumors also exhibited the highest levels of extracellular IL-33, which was associated with enhanced IFN-ɣ production and upregulation of the IL-33 receptor ST2 in tumor-infiltrating T cells. Although CD8+ T cells contributed to tumor control, their depletion did not abolish the differential growth between *FL*- and *Ala*-expressing tumors, suggesting that additional cell populations contribute to tumor control. However, genetic ablation of *St2* abolished the differential tumor growth associated with the IL-33 isoforms, demonstrating that host ST2 signaling is required for isoform-dependent tumor control. Taken together, our findings identify extracellular cytokine availability as a key determinant of the antitumor activity of tumor-derived IL-33 and support a model in which IL-33 promotes ST2-dependent effector T-cell function. Furthermore, our findings highlight intracellular localization versus extracellular release of IL-33 as potential determinants of the context-dependent effects of the IL-33/ST2 axis in cancer.

## Introduction

The tumor microenvironment (TME) is a complex system in which malignant cells interact with immune, stromal and vascular cells to determine tumor progression versus immune response. Cytokines within the TME modulate these interactions by regulating immune-cell recruitment, activation, differentiation and function. Among these cytokines, interleukin-33 (IL-33) has emerged as an important regulator of tumor immunity, with both tumor-promoting and tumor-suppressing effects being reported, depending on the tumor type, cellular context and inflammatory environment^1–3^.

IL-33 belongs to the IL-1 family of cytokines and shares structural similarities with other members of this family, as well as signaling through a heterodimer receptor, composed of the shared IL-1 family receptor IL1RAcP (encoded by *IL1RAP*) and ST2 (encoded by *IL33R,* representing the IL-33-specific moiety)^4,5^. IL-33 is constitutively expressed in the nucleus of cells lining the tissue, such as epithelial, endothelial, and fibroblast-like cells^6,7^. There, IL-33 is present as a precursor form, also known as full-length IL-33.

Upon cell stress, damage, or necrosis, IL-33 is released into the extracellular space, where it can undergo proteolytic processing to generate mature IL-33, which subsequently binds to ST2-expressing immune cells^5^. IL-33/ST2 signaling has been reported to modulate both innate and adaptive immune responses by acting on multiple immune cell subsets^8,9^, including mast cells^10^, type 2 innate lymphoid cells (ILC2s), natural killer (NK) cells, CD8+ T cells^11,12^, and distinct CD4+ T cells subsets, including T regulatory cells (Tregs)^13^.

Although nuclear IL-33 can bind to chromatin through the acidic pockets of the H2A-H2B histone dimer^14,15^, whether this interaction plays a role in transcriptional regulation remains unclear. Several recent studies have described ST2-independent functions of IL-33 that are associated with its nuclear localization. In the nucleus, IL-33 can interact with transcription factors such as RUNX2^16^ and GATA3^17^, thereby modulating signaling pathways, including TGF-β/SMAD^16,18^ and HIF-1α^19^. Moreover, Tregs and myeloid cells have been reported to express IL-33 and thereby modulate immune response^20–22^. However, immune cells are not generally considered as *bona fide* IL-33 producing cells^23^. The proposed nuclear function of IL-33 is not entirely established and remains controversial, as genetic and transcriptional studies have not identified convincing transcriptional consequences of endogenous nuclear IL-33^23^. Apart from a potential nuclear function, localization (or sequestration) in the nucleus controls IL-33’s cytokine activity by preventing excess release^6^. Proteolytic processing of full-length IL-33 by extracellular inflammatory proteases^6^ or caspases^24^ may be critical for regulating immune responses^6^. This is particularly relevant given that mature IL-33 exhibits substantially greater biological activity than its full-length counterpart^25,26^.

This distinction between full-length and mature IL-33 may be particularly relevant in cancer, where IL-33 can exert both anti- and pro-tumorigenic effects, potentially depending on the nature and composition of IL-33-responsive, ST2-expressing cell populations within the TME^1–3^. By enhancing the activation and accumulation of CD8+ T cells and NK cells in the TME^2,3^, IL-33 can on the one hand promote anti-tumor immunity^27^. On the other hand, IL-33/ST2 signaling may promote the proliferation and recruitment of tumor-infiltrating Tregs, or activate myeloid-derived suppressor cells, which enhances immune suppression and tumor progression^2,3,28^. Illustrating these dual, context-dependent effects, MC38 murine adenocarcinoma cells overexpressing IL-33 were reported to exhibit increased tumor growth compared with control cells^29^. Conversely, intraperitoneal administration of recombinant IL-33 suppressed MC38 tumor growth and increased tumor-infiltrating T cell numbers^27^. Thus, the net effect of IL-33/ST2 signaling on tumors may depend on the specific ST2+ immune cell populations present within the TME and how they respond to IL-33.

Here, we developed an experimental system in which MC38 cells express distinct IL-33 isoforms with defined nuclear retention and extracellular availability, allowing us to investigate the largely unexplored contribution of individual IL-33 isoforms to tumorigenesis. Specifically, we generated polyclonal MC38 cell lines expressing full-length, immature IL-33; a full-length isoform of IL-33 with mutations in its chromatin binding-domain; or truncated, mature IL-33 lacking its nuclear functional domains. By comparing these isoforms *in vitro* and *in vivo*, we found that increased extracellular availability of IL-33 was associated with stronger control of tumor growth, enhanced IFN-ɣ production of tumor-infiltrating T cells, and increased ST2 expression on immune cells. Although CD8+ T cells contributed to tumor growth inhibition, their depletion did not fully abrogate the differences in tumor progression observed between the cell lines, suggesting involvement of additional immune populations. Importantly, genetic ablation of *St2* abolished the differences in tumor development between the IL-33 isoforms, demonstrating that the anti-tumor effects of tumor-derived IL-33 require host ST2 signaling. Collectively, these findings establish extracellular availability and cytokine function as key determinants of IL-33 mediated tumor control and provide insight into how the localization of tumor-derived IL-33 shapes the immune response within the TME.

## Results

### Generation and molecular characterization of MC38 cells expressing distinct IL-33 isoforms

To investigate the intracellular and extracellular functions of IL-33 in tumor epithelial cells, we generated lentiviral vectors encoding IL-33 isoforms with distinct properties and used them to establish different polyclonal MC38 mouse adenocarcinoma cell lines (MC38-*Il33*) (Figure 1A-B). The individual IL-33 isoforms were designed to selectively retain or disrupt specific functional domains of IL-33 (Figure 1C), thereby enabling independent investigation of its nuclear versus extracellular activity. The first construct included a full length (*FL*) isoform of IL-33, encoding the canonical IL-33 protein and including both its intracellular/nuclear and extracellular functions. As a second isoform, we generated an alanine-mutated IL-33 isoform (*Ala*) carrying a triple mutation in the region of murine *Il33* encompassing residues L49-S51 (Leu49Ala, Arg50Ala and Ser51Ala), since mutation of these residues to alanine has been reported to abrogate chromatin association without affecting the nuclear localization of IL-33^15^. However, the *Ala* construct still includes the region encompassing amino acids 66-110, which has been reported to interact with nuclear factor kB (NF-kB)^30^. The third construct encoded for a truncated IL-33 isoform (*Trunc*), corresponding to the processed, mature form of IL-33. This isoforms lacks all regions required for IL-33’s nuclear functions, while retaining the bioactive C-terminal domain of the cytokine^5,25^. It therefore enables a focused assessment of the extracellular activity of IL-33 in the absence of its nuclear domains. Further, the mature form of IL-33 displays significantly enhanced biological activity relative to its full-length counterpart^25,26^.

**Figure 1:**
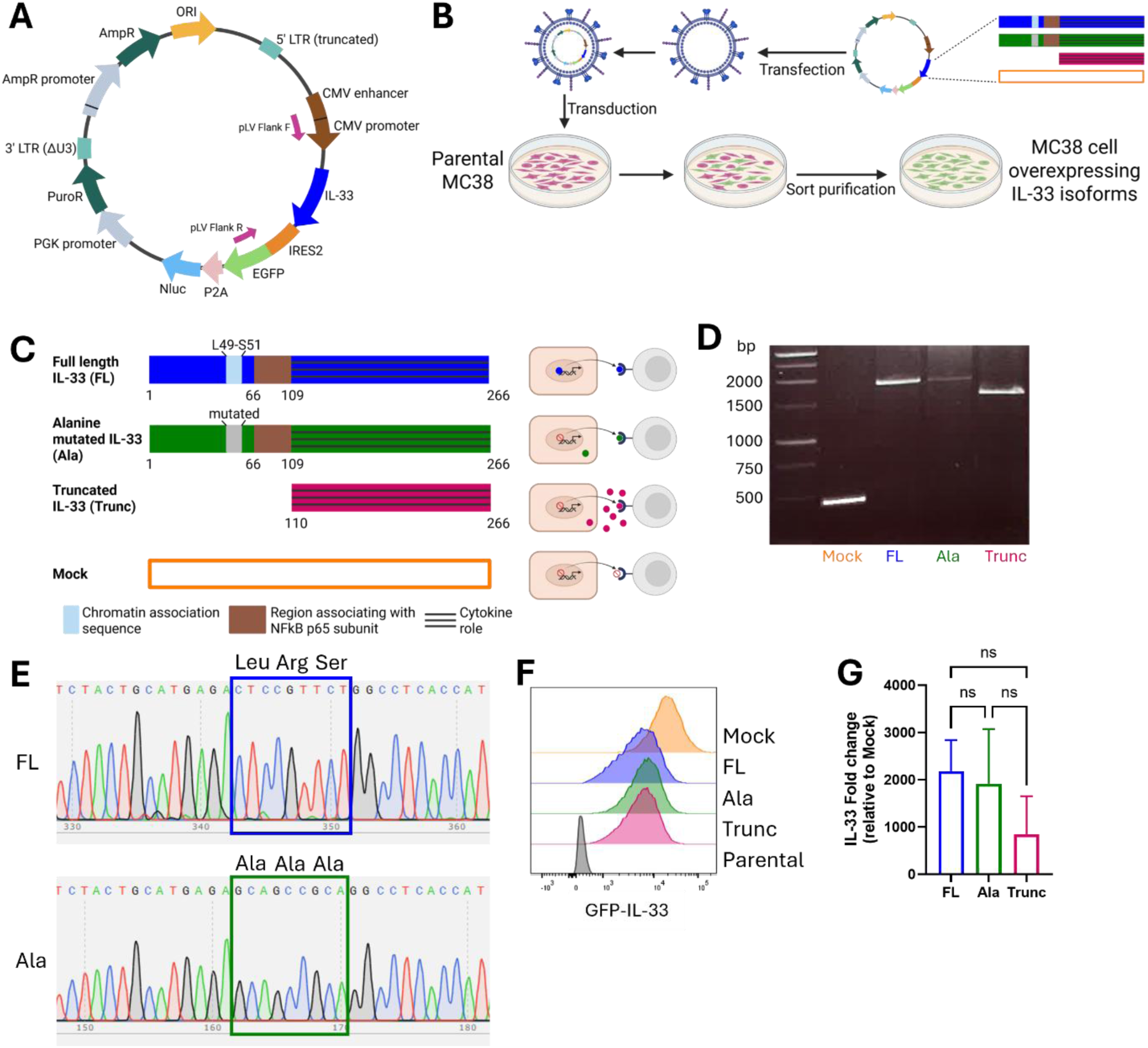
Generation of MC38 cell lines overexpressing different IL-33 isoforms. (A) Backbone of the vector plasmid, exemplary for FL IL-33. Primers used for PCR detection are indicated by pink arrows. (B) Schematic overview of the generation of the different MC38-Il33 cell lines. MC38-Il33 cells were transduced using a lentiviral system and subsequently selected using puromycin and sort purifications. (C) Overview of the different IL-33 isoforms: Full length (FL), Alanine mutated (Ala), Truncated (Trunc) and Mock control. (D) PCR detection of the different transgenic IL-33 isoforms. Expected sizes: FL and Ala: 18S2 base pairs (bp, Trunc: 1571 bp, Mock: 44S bp). Representative image of 2 independent experiments is shown. (E) Alignment of the FL and Ala sequences after Sanger sequencing. Sequencing confirmed the triple alanine mutation (Leu4SAla, Arg50Ala, Ser51Ala) within the chromatin-binding motif of the Ala cell line. (F) GFP expression of the cell lines after the sort purification. (G) Expression levels of Il33 of the cell lines normalized to the Mock. Data were conducted in 4 biological replicates. 1-way ANOVA with Tukey’s multiple comparisons test was used (ns).

As a cell line, we selected the MC38 mouse adenocarcinoma cell line over the polyploid mouse carcinoma cell line CMT-93, since endogenous *Il33* transcripts and IL-33 protein were not detectable in parental MC38 compared to CMT-93 cells (Supplementary Figure 1A-D). MC38 cells have been widely used as an *in vivo* model of tumor showing robust growth following transplantation^29,31,32^. In contrast to the lack of IL-33 expression, MC38 expressed *St2* transcripts, which were absent in CMT-93 cells (Supplementary Figure 1E). Further analysis using *St2* isoform-specific primers suggested predominant transcriptional expression of the soluble ST2 (sST2) isoform in MC38 cells (Supplementary Figure 1D).

Genomic integration of the IL-33 transgene was confirmed by PCR, with amplicon sizes consistent with those expected for each construct (Figure 1D). Presence of the targeted triple mutation (Leu49Ala, Arg50Ala, Ser51Ala) within the chromatin-binding motif of the *Ala* MC38 cell line was verified by Sanger sequencing (Figure 1E). Flow cytometric sorting yielded uniformly GFP-positive populations (100%) across all transduced MC38 cell lines (Figure 1F), and *Il33* expression levels were similar across the different MC38-*Il33* lines, supporting their suitability for subsequent functional comparisons (Figure 1G).

In summary, we generated three polyclonal MC38 cell lines overexpressing IL-33 isoforms with distinct nuclear and extracellular properties, providing a tool to dissect the contribution of the different IL-33 functions to tumor growth.

### *In vitro* characterization of MC38-*Il33* cell lines

We next phenotypically characterized the individual cell lines expressing distinct IL-33 isoforms and assess their respective IL-33 transgene expression levels. Immunohistochemistry confirmed the expected differences in subcellular localization (Figure 2A-E). *FL* IL-33 was primarily detected in the nucleus, whereas the *Ala* and *Trunc* IL-33 isoforms predominantly localized in the cytoplasm. In contrast, both parental and *Mock* vector-transduced MC38 cells showed only marginal IL-33 staining, detectable in 3-5% of cells (Supplementary Figure 2A-B), potentially reflecting specific background staining or low endogenous IL-33 expression.

**Figure 2:**
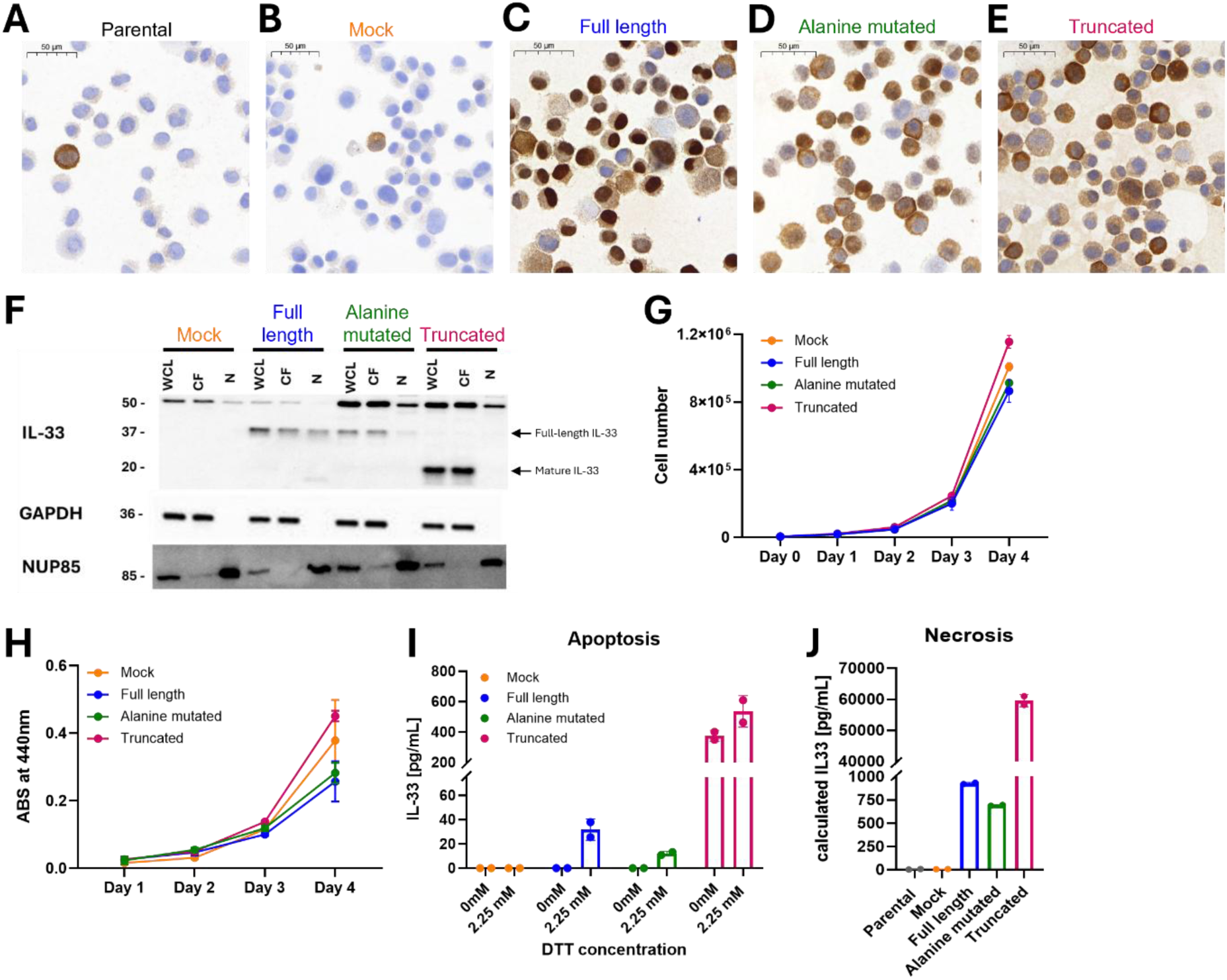
In vitro characterization of MC38-Il33 cell lines. Immunohistochemical staining for IL-33 (brown) in MC38-Il33 cell lines using the cytospin method. (A) Parental, unmodified MC38 cells, (B) Mock, (C) Full length, (D) Alanine mutated and (E) Truncated MC38-Il33 cells. (F) Western blot-based detection and localization of the IL-33 protein. IL-33 was analyzed in whole-cell lysate (WCL), cytosolic fraction (CF) and nuclear fraction (N) of MC38-Il33 cell lines. GAPDH and NUP85 were used as controls for the cytosolic and nuclear fractions, respectively. GAPDH and NUP85 were used as cytoplasmic and nuclear fractionation controls, respectively. Expected sizes of full-length (34 kDa) and truncated (1S kDa) IL-33 are indicated. (G) In vitro proliferation of MC38-Il33 cell lines. Cells were counted with a CASY counter. (H) Baseline corrected absorbance of WST cell proliferation assay. (I) 2.25 mM DTT-induced apoptosis (**Fehler! Verweisquelle konnte nicht gefunden werden.**C) and ELISA-based detection of IL-33 in cell supernatants. Medium (0mM) or 2.25mM DTT were added to the indicated cell lines for 24h. (J) Necrosis-driven IL-33 release of indicated MC38 cell supernatants upon freeze-thaw cycles measured by ELISA. Truncated IL-33 levels were calculated using the spike-and-recovery method. (G-H) Representative of 2 independent experiments is shown, N=3 technical triplicates. 1-way AONVA with Tukey’s multiple comparisons test was used (ns). (I-D) Representative data from 1 experiment is shown, N=2.

These findings were further validated by western blot analysis of whole-cell lysate (WCL), cytosolic fraction (CF) and nuclear fraction (N) of the different MC38-*Il33* cell lines (Figure 2F). In line with the histology data, IL-33 protein was not detected in *Mock* cells, while the individual IL-33 isoforms migrated at the expected molecular weight – 34 kDa for full-length IL-33 and 19 kDa for truncated, mature IL-33^25^. *FL* IL-33 isoform was detected in all three cellular fractions. In contrast, the *Ala* IL-33 and *Trunc* IL-33 variants were absent from the nuclear fraction. An additional, unspecific, band at approximately 50 kDa was detected in all cell lines, suggesting unspecific binding to a MC38 cell-specific antigen. Overall, both immunohistochemistry and western blot analysis confirmed the expression and the expected localization of the individual IL-33 isoforms in the MC38-*Il33* cell lines.

To assess potential effects on cell proliferation, the different MC38-*Il33* cell lines were monitored over time. Cell counting over a four-day period revealed comparable growth rates across all cell lines (Figure 2G), a finding further corroborated by a WST-based proliferation assay (Figure 2H). Our findings indicated that the individual isoforms did not alter *in vitro* growth kinetics, making them also suitable for comparison in an *in vivo* setting.

Finally, extracellular IL-33 release was examined. Relative to the *FL* and *Ala* IL-33 isoform, the *Trunc* variant was detected at substantially higher levels in MC38 cell line supernatant. This was the case under basal conditions, upon apoptosis induction using dithiothreitol (DTT) (Figure 2I), but also following freeze-thaw-induced necrosis (Figure 2J) – which were tested since IL-33 had been reported to be released in response to apoptotic cell death^6^ or necrotic cellular stress^33^. Both apoptosis and, more prominently, necrosis promoted the extracellular release of the *FL* and *Ala* IL-33 isoforms, which were otherwise undetectable in the supernatants of unstressed cells.

In sum, these results demonstrated preferential extracellular release of the mature *Trunc* IL-33 isoform compared to the *FL* IL-33 and *Ala* IL-33 counterparts, suggesting that impaired nuclear retention may underlie this phenotype.

### IL-33 overexpression suppresses MC38 tumor growth *in vivo*

Having established that the MC38-*Il33* cell lines exhibited comparable proliferation rates *in vitro*, we next investigated whether the individual IL-33 isoforms may alter tumor development *in vivo*. Following injection into both flanks of wild-type (WT) mice, *Mock* cells gave rise to pronounced tumor outgrowth, whereas tumor volumes were generally lower in animals injected with IL-33-isoform-expressing MC38 cells (Figure 3B-C and Supplementary Figure 3). Among the individual IL-33 isoforms, expression of *Trunc* IL-33 resulted in the greatest reduction in tumor growth. This effect may reflect the previously reported higher biological activity of mature IL-33^25^, its greater extracellular release (Figure 2I), or a combination of both. Tumors expressing *Ala* IL-33 also exhibited reduced growth compared to *FL*-IL-33-expressing counterparts.

**Figure 3:**
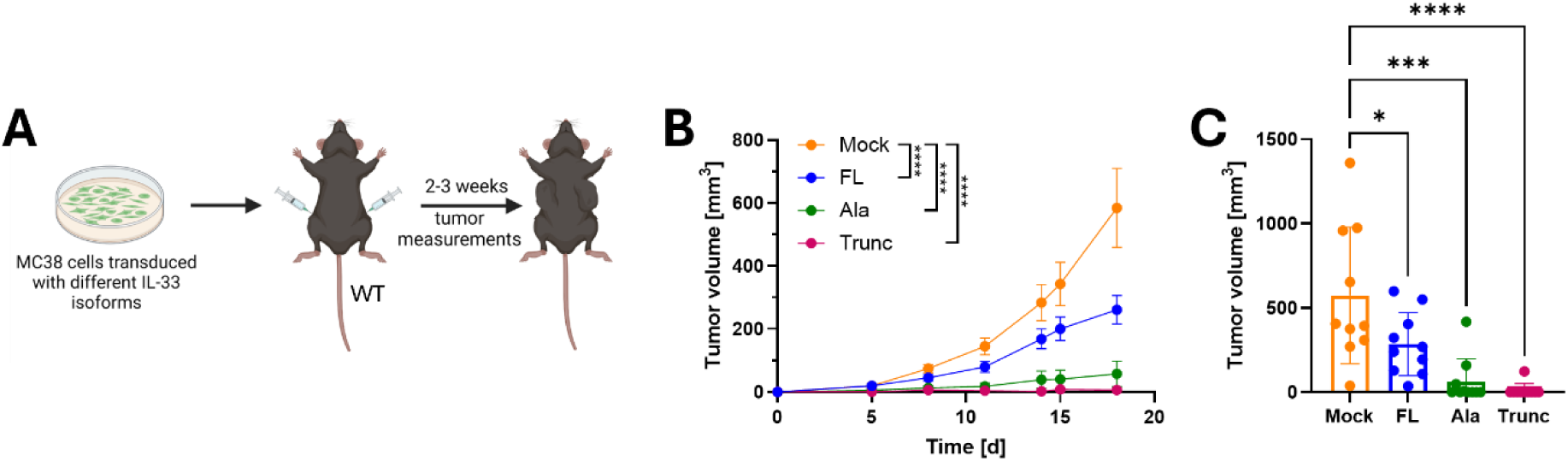
IL-33 overexpression decreases MC38 tumor growth. (A) Schematic overview of the experiment. MC38 cell lines overexpressing different IL-33 isoforms were subcutaneously injected into both flanks of WT mice. Tumor volume was measured every 2-3 days with a caliper. (B) Tumor growth of the different IL-33 isoforms over time. (C) Tumor volume at the endpoint (d18). Data pooled from 2 independent experiments, N=10 mice per group. (B) 2-way ANOVA and (C) 1-way ANOVA with Tukey’s multiple comparisons test were used. *: P < 0.05; **: P < 0.01; ***: P < 0.001; and ****: P < 0.0001. (A) Figure generated with Biorender.com.

Together, these findings indicated that the individual IL-33 isoforms differentially affected tumor development *in vivo*, with nuclear retention of IL-33 being associated with enhanced tumorigenesis. Nevertheless, IL-33 overexpression overall resulted in reduced tumor growth.

### IL-33 promotes CD8+ T cell- and IFN-ɣ-associated tumor control

To assess potential immune mechanisms underlying the differential growth of the MC38-*Il33* cell lines in recipient mice, we analyzed tumor-infiltrating immune cells in *FL* and *Ala* IL-33-expressing tumors. We focused on *FL* and *Ala* IL-33-expressing tumors, because these isoforms differed in their nuclear localization, as observed in our western blot data (Figure 2F). *Trunc* IL-33 MC38 cells were not used, because their engraftment was inconsistent, with some mice failing to develop tumors, thereby preventing a reliable analysis of tumor-infiltrating immune cells.

With the exception of CD8+ T cells, which were more abundant in *Ala* IL-33-expressing tumors, the frequencies of the other tumor-infiltrating immune cell subsets did not differ between the experimental groups (Supplementary Figure 4A-C). Based on these findings and given the established role for CD8+ T cells in controlling MC38 tumor growth^27,34^, we next investigated their contribution to IL-33 isoform-dependent MC38 tumor control.

*FL* and *Ala* IL-33 cells were subcutaneously injected into the flanks of WT mice, followed by treatment with either anti-CD8-depleting antibody or isotype control (Figure 4A). CD8+ T cell depletion led to increased tumor growth in both groups but it did not abrogate the difference in tumor size between the two cell lines (Figure 4B and Supplementary Figure 4D-E). Instead, the difference in tumor growth between *FL* and *Ala* IL-33 tumors was maintained following CD8+ T cells depletion, which was also reflected in the final tumor weights (Figure 4C). Moreover, tumor weight negatively correlated with the proportion of CD45+ tumor-infiltrating immune cells (Figure 4D). These findings confirm the contribution of CD8+ T cells to MC38 tumor control, while also suggesting that additional immune or non-immune cell mechanisms contribute to the differential growth of *FL* versus *Ala* IL-33-expressing tumors.

**Figure 4:**
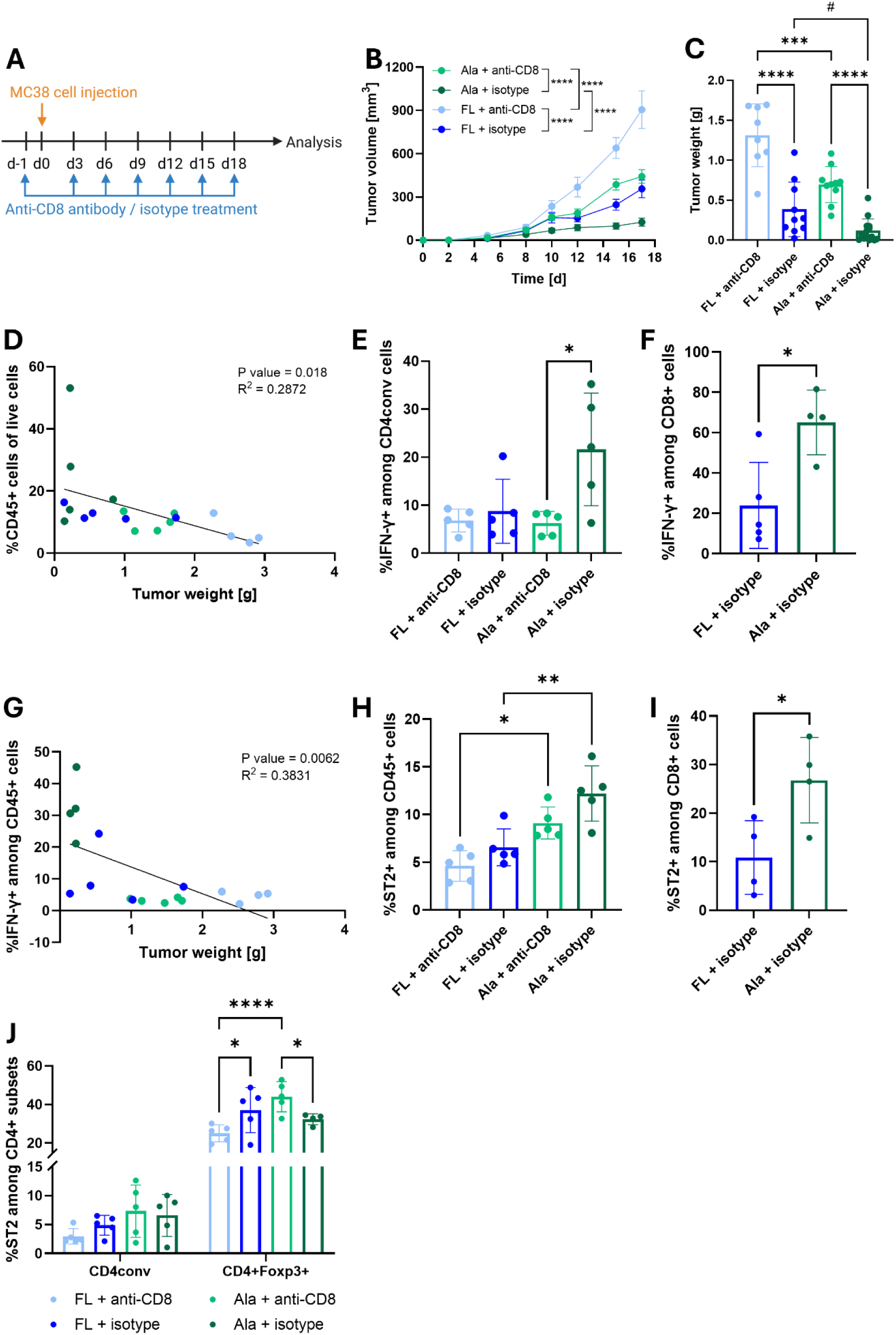
IL-33-dependent tumor growth is in part controlled by CD8+ T cells. (A) Schematic overview of the experimental setup. WT mice were subcutaneously injected into both flanks with FL or Ala MC38 cells in combination with either depleting anti-CD8 antibody or isotype control treatment every 3 days. At the endpoint, tumors were excised and digested for analysis of tumor-infiltrating immune cells. (B) Tumor growth of the indicated groups over time (C) Tumor weight at the endpoint (d1S). # indicates significance determined by unpaired t test between FL+isotype and Ala+isotype. (D) Correlation of CD45+ cells versus tumor weight at the endpoint. IFN-ɣ+ cells among (E) conventional CD4+ T cells (CD4conv) and (F) CD8+ T cells. (G) Correlation of IFN-ɣ+CD45+ cells versus tumor weight at the endpoint. ST2+ cells among (H) CD45+ immune cells, (I) CD8+ T cells (J) CD4conv and CD4+Foxp3+ cells and. (B-J) Data from 1 experiment, N=5-7 mice per (B) 2-way ANOVA with Tukey’s multiple comparisons test. (C, E, H, J) 1-way ANOVA with Tukey’s multiple comparisons test, (C#, F, I) Unpaired t test with Welch’s correction. (D-G) Linear regression analysis. *: P < 0.05; **: P < 0.01; ***: P < 0.001; and ****: P < 0.0001. group. (A) generated with Biorender.com.

As previous studies have implicated IFN-ɣ in MC38 tumor control^27,34,35^, we next assessed IFN-ɣ production in tumor-infiltrating lymphocytes. Frequency of IFN-ɣ-producing cells was significantly increased in conventional CD4+ T cells (CD4conv) in *Ala* IL-33 tumors compared with the other groups (Figure 4E, experimental group *isotype*). Similarly, CD8+ T cells from *Ala* IL-33 tumors exhibited an increased frequency of IFN-ɣ+ cells (Figure 4F). Consistent with these results, the frequency of IFN-ɣ+ cells among total CD45+ immune cells was increased in *Ala* IL-33 tumors and negatively correlated with tumor weight (Figure 4G and Supplementary Figure 4F). Notably, CD8+ T cell depletion did not affect the proportion of IFN-ɣ+ cells among CD4conv T cells in *FL* IL-33 tumors.

As a next step, we investigated ST2 expression to further assess the relationship between the extracellular availability of IL-33 and immune cell responses. ST2 was specifically expressed on immune cells within tumors, but not detected on non-immune cells (Supplementary Figure 4G). Overall, the proportion of ST2+ cells among CD45+ immune cells was higher in *Ala* IL33 tumors than in *FL* IL-33 counterparts, irrespective of CD8+ T cell depletion (Figure 4H). Although myeloid cells, particularly CD11b+CD11c+ cells, constituted the largest proportion of tumor-infiltrating immune cells, they did not express ST2 (Supplementary Figure 4H). Further analysis revealed that ST2 expression was restricted to T cells (Supplementary Figure 4I-N). In the isotype control-treated group, *Ala* IL-33 tumors exhibited a higher frequency of ST2-expressing CD8+ T cells than *FL* IL-33 tumors, with a similar trend towards increased per-cell ST2 expression on CD8+ T cells in *Ala* IL-33 tumors (Figure 4I and Supplementary Figure 4I-J). In all groups, frequency of ST2+ cells was low among CD4conv T cells, but substantially higher among CD4+Foxp3+ Tregs (Figure 4J and Supplementary Figure 4K-N). While frequency of ST2+ Tregs was similarly high in *FL* versus *Ala* IL-33 tumors, CD8+ T cell depletion resulted in a modest but statistically significant change in the frequency of ST2-expressing Tregs in *Ala* compared with *FL* IL-33 tumors.

Together, these results suggest that reduced nuclear retention of IL-33 in *Ala* IL-33 MC38 cells was associated with enhanced ST2 expression on tumor-infiltrating T cells and increased IFN-ɣ production, particularly by CD8+ T cells. The persistence of differences in tumor growth following CD8+ T cell depletion in *FL* versus *Ala* IL-33 tumor groups suggests that additional ST2-expressing populations, possibly Tregs, may also control tumor growth in an IL-33-dependent manner.

### ST2 is required for IL-33-mediated tumor control

Having identified increased ST2 expression on tumor-infiltrating T cell subsets in IL-33-overexpressing MC38 tumors, we next investigated whether ST2 was required for the differential control of MC38-*Il33* tumors. For this, the different MC38-*Il33* cell lines were injected subcutaneously into both flanks of *St2^-/-^* mice (Figure 5A). In the global absence of *St2* in host mice, tumors expressing the different IL-33 isoforms exhibited comparable tumor growth (Figure 5B-C and Supplementary Figure 5).

**Figure 5:**
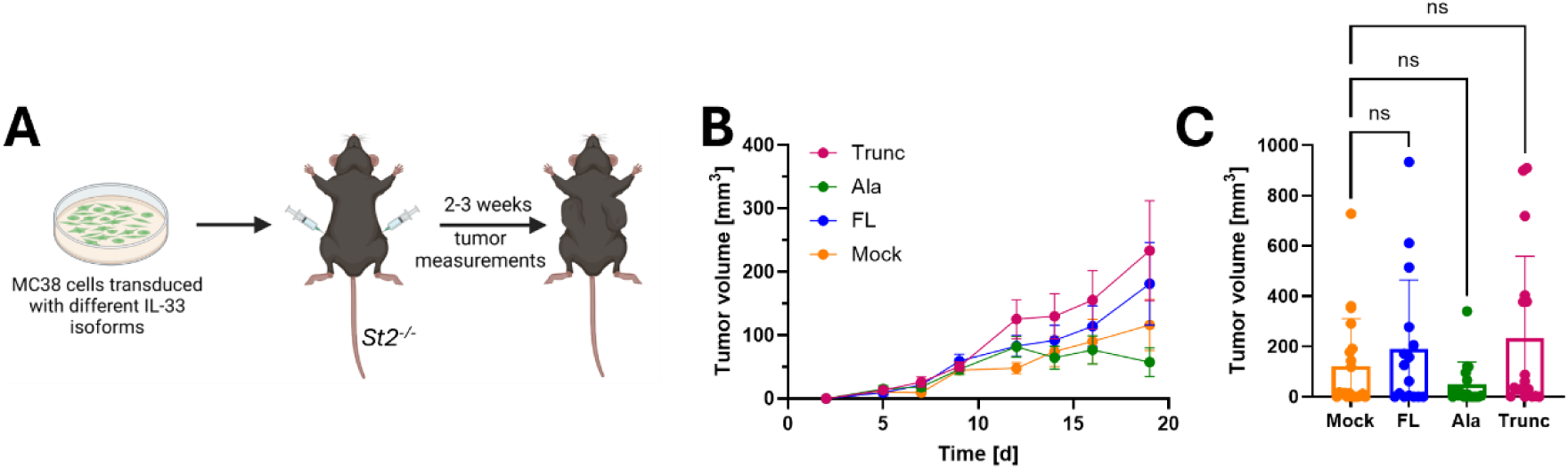
Ablation of St2 abrogates the anti-tumor effect of IL-33. (A) Schematic overview of the experiment. MC38 cell lines overexpressing different IL-33 isoforms were subcutaneously injected into both flanks of St2^-/-^ mice. Tumor volume was measured every 2-3 days with a caliper. (B) Tumor growth of the different IL-33 isoforms over time. (C) Tumor volume at the endpoint (d1S). Data pooled from 2 independent experiments, N=10-12 mice per group. (B) 2-way ANOVA and (C) 1-way ANOVA with Tukey’s multiple comparisons test were used (ns). (A-C) Figure generated with Biorender.com.

Together, these findings indicate that genetic ablation of *St2* abolished the differences in tumor growth observed between the IL-33 isoform-expressing MC38 cell lines in WT mice. This suggests that ST2-expressing immune cells within the TME modulate tumor growth depending on extracellular IL-33 availability.

## Discussion

IL-33 is a pleiotropic cytokine with both intracellular nuclear functions and extracellular cytokine activities. In cancer, the IL-33/ST2 axis has been associated with both tumor-promoting and tumor-suppressive effects, depending on the tumor type, cellular source of IL-33 and the composition of the TME^6,36^. In the present study, we addressed this complexity by generating MC38 cells expressing distinct IL-33 isoforms that differ in their capacity for nuclear retention and extracellular availability. This approach allowed us to distinguish the effects associated with nuclear IL-33 from those associated with extracellular IL-33 signaling via the ST2 receptor. We found that tumor-cell-derived IL-33 suppressed MC38 tumor growth *in vivo*, with stronger tumor control in recipient mice being associated with decreased nuclear localization capacity and/or increased extracellular availability of IL-33 isoforms. Importantly, the differential tumor growth observed between the IL-33 isoforms was abolished in *St2^-/-^* mice, indicating that ST2 signaling is required for tumor control in our model. Together, these findings suggest that the biological activity of tumor-derived IL-33 is determined, at least in part, by the extent to which IL-33 is retained intracellular versus made available for extracellular ST2-dependent signaling.

The distinct localization of the IL-33 isoforms in our MC38 cell lines provides a mechanistic basis for their differential activity. Full-length IL-33 is controlled by its localization (or sequestration) in the nucleus, which prevents excess release^6^. Processing of full-length IL-33 into its more biologically active mature form by extracellular inflammatory proteases^6^ or caspases^24^ is critical in regulating immune responses^6^. Consistent with this, *FL* IL-33 was predominantly localized in the nucleus of MC38 cells, as expected^6^. In contrast, disruption of the chromatin-binding motif in the *Ala* isoform resulted in loss of detectable nuclear IL-33, as described previously^15^. The *Trunc* IL-33 isoform, which lacks the nuclear functional domains of the cytokine, was predominantly cytoplasmic in MC38 cells – in line with a previous observation that mice overexpressing truncated IL-33 exhibited elevated serum levels of this cytokine, suggesting that removal of its nuclear retention domains promotes the constitutive release of IL-33^37^.

The differences in tumor growth *in vivo* were unlikely to result from intrinsic effects of the individual isoforms, as expression of the different constructs did not substantially affect MC38 cell proliferation *in vitro*. Instead, tumor growth inversely correlated with extracellular IL-33 availability. This observation is consistent with the established function of IL-33 as an alarmin that is released following cellular stress and subsequently activates ST2-expressing immune cells^6,36^.

Notably, *Trunc* IL-33 was detectable extracellularly even in the absence of experimentally induced cell death, and it was released at particularly high levels following apoptotic or necrotic stress. The *Trunc* IL-33 isoform, being already in its mature form, not only was more readily released extracellularly, but it is also expected to exhibit greater intrinsic biological activity than the full-length protein, without requiring further proteolytic processing^25,26^. This may have further contributed to the more pronounced reduction in tumor growth observed in mice bearing *Trunc* IL-33-expressing tumors.

Although we did not determine the levels of full-length and mature IL-33 within the tumors, our *in vitro* findings suggest isoform-specific differences in their release. The intermediate *in vivo* tumor growth phenotype observed with *Ala* IL-33 further supports a role for chromatin association in regulating IL-33 availability. Although *Ala* and *FL* IL-33 did not differ in their release following experimentally-induced cell death *in vitro*, disruption of the chromatin-binding motif resulted in loss of detectable nuclear IL-33 as detected by western blot analysis and increased tumor control in recipient mice. These observations align with previous evidence that chromatin binding promotes nuclear retention of IL-33 and limits its extracellular release^38^. Together, these findings extend the concept of IL-33 localization beyond a simple distinction between nuclear and extracellular IL-33 and suggest that the degree of intracellular retention can influence the amount of tumor-derived IL-33 available for ST2-dependent signaling.

An alternative explanation for the differential tumor growth is that nuclear IL-33 potentially exerts cell-intrinsic effects in MC38 cells. Several studies have proposed that nuclear IL-33 can regulate cellular functions independently of ST2. For example, *Park et al.* demonstrated that full-length, but not mature, IL-33 activates nuclear TGF-β/SMAD signaling, thereby promoting epithelial cell proliferation and skin carcinogenesis^16^. Nuclear IL-33 has also been shown to suppress GATA-mediated soluble ST2 (sST2) expression in colorectal cancer cells under hypoxic conditions, with restoration of sST2 expression inhibiting heterotopic tumor growth and lung metastasis^17^. In addition, nuclear IL-33 has been proposed to suppress the activation of HIF-1α in intestinal epithelial cells^19^. These observations raise the possibility that *FL* IL-33 could exert nuclear, cell-intrinsic effects that are lost following disruption of chromatin association in the *Ala* IL-33 isoform. However, the proposed nuclear functions of IL-33 remain incompletely defined, as genetic and transcriptional studies have not yet provided convincing evidence for substantial transcriptional effects of endogenous nuclear IL-33^23^. Together with the comparable *in vitro* proliferation rates of the different MC38 cell lines, which argue against cell-intrinsic effects of the individual IL-33 isoforms, the ablation of differential tumor growth in *St2^-/-^*mice further supports a requirement for extracellular ST2 signaling in driving the distinct *in vivo* growth phenotypes observed in WT recipients. Nevertheless, a contribution of nuclear IL-33 to the *FL* phenotype cannot be excluded and would require direct analysis of IL-33-dependent transcriptional programs in the MC38 cells.

ST2 expression within the tumor lesions was restricted to immune cells, while we only investigated and detected *sSt2* expression on MC38 cells *in vitro*. Although the potential contribution of tumor-derived sST2 was not investigated further, our findings support a model in which tumor-derived IL-33 primarily exerts its effects through ST2-expressing immune cells. This interpretation is consistent with previous studies showing that tumor-derived or exogenously administered IL-33 can promote CD8+ T cell accumulation and effector function, including IFN-ɣ production, and contribute to anti-tumor responses in MC38 tumors^27,34,39^.

Our antibody-mediated depletion experiments further indicated that CD8+ T cells contribute to, but do not fully account for, the differential tumor control. CD8+ T cell depletion increased tumor growth in both *FL* and *Ala* IL33-expressing tumors, confirming an important role for CD8+ T cells in MC38 tumor control. However, the difference between *FL* and *Ala* IL-33 tumors persisted after this cell depletion. This points to the involvement of additional immune cell populations in IL-33-dependent tumor control. In line with this interpretation, the frequency of IFN-ɣ-producing cells was increased not only in CD8+ T cells but also in conventional CD4+ T cells in *Ala* IL-33 tumors. In addition to T cells, we also detected IFN-ɣ production in NK and NKT cells, with the highest frequencies observed in *Ala* IL-33 tumors (data not shown). Although previous studies have not specifically implicated IFN-ɣ-producing NK and NKT cells in IL-33-dependent MC38 tumor control^27,34,39^, these populations may contribute to the enhanced tumor control observed in our model. Increased IFN-ɣ production may indeed result from direct activation of ST2-expressing T cells or from indirect effects of IL-33 on other ST2-negative immune populations. Additional CD8+ T cell effector mechanisms, including the production of granzyme B and perforin and the expression of Fas ligand, have been implicated in tumor control^40–42^, however, their contribution to the phenotype observed in our study remains to be determined.

The composition of the ST2-expressing immune compartment may be an important determinant of the outcome of IL-33 signaling. In our model with MC38 cell lines, ST2 was highly expressed on Tregs and CD8+ T cells, but only marginally on CD4conv T cells. This pattern is consistent with previous reports showing that ST2 can be particularly prominent on activated or effector T cells and on tumor-infiltrating Tregs^28,34,39,43^. Supporting this notion, *Gao et al.* reported enhanced tumor elimination when IL-33 overexpression was combined with Treg depletion in a B16 melanoma model^39^. Together, these studies support the possibility that Tregs modulate IL-33-regulated anti-tumor immunity. Together, these studies support the possibility that Tregs modulate IL-33-regulated anti-tumor immunity. However, we did not investigate the suppressive functions of Tregs in our models, and further studies are needed to better define their contribution to the observed phenotype.

The importance of the cellular context of ST2 signaling is further highlighted by studies indicating opposing effects of *St2* deficiency on tumor growth. *Son et al.* reported that *St2* deficiency enhanced anti-tumor immunity in a murine lung tumor model^43^. In their study, loss of *St2* reduced the accumulation of tumor-infiltrating Tregs, while increasing the frequency of CD8+ T cells and their IFN-ɣ production. Similarly, *Van der Jeught et al.* reported reduced tumor growth in the MC38 tumor model upon genetic ablation of *St2*, which was associated with an enhanced CD8+ T cell-mediated anti-tumor response^44^. However, the authors did not detect ST2 expression on CD8+ T cells in their model. Instead, their data implicated ST2+ tumor-associated macrophages and ST2+ Tregs as key immune populations through which IL-33 modulated tumorigenesis. A role for tumor-associated macrophages in IL-33-dependent MC38 tumor control was, until now, only described by *Zhang et al.*, although their ST2 expression was not assessed^29^. These findings appear to be at odds with our observation that ST2 is required for the tumor-suppressive effects of cancer cell-derived IL-33. One possible rationale for these apparently opposing findings is that the consequences of ST2 signaling may differ depending on the immune cell population in which it is activated. In support of this concept, an alternative second promoter for the *St2* gene has recently been identified^45^, in addition to the previously characterized regulatory elements associated with *St2* expression in type 2 immune cells and Tregs^12^, this second promoter regulates *St2* expression in type 1 immune responses and contributes to antiviral cytotoxic T cell function by supporting T cell receptor stimulation^45^.

The amount and spatial availability of IL-33 may represent an additional determinant of this balance, depending on whether it is present locally within the TME, in the surrounding tissues, or systemically. In our model, the *Trunc* IL-33 isoform increased extracellular IL-33 availability and was associated with enhanced tumor control. One explanation for this observation is that higher local concentrations of IL-33 may favor activation of ST2-expressing CD8+ T cells, thereby promoting the anti-tumor response. Conversely, lower levels of IL-33 might be more readily captured by ST2+ Tregs, potentially promoting their accumulation and suppressive function. A similar principle has been described for IL-2, whereby Tregs efficiently limit its availability through high CD25 expression, thereby restricting CD8+ T cell expansion and effector differentiation^46,47^.

However, recent findings suggest that the relationship between IL-33 availability and Treg function may be more complex than a simple concentration-dependent competition between CD8+ T cells and Tregs. *Liu et al*. recently reported that systemic IL-33 treatment increased the abundance of ST2+ Tregs, while nevertheless suppressing tumor growth in several mouse tumor models, including MC38 tumors^48^. Importantly, this anti-tumor effect was dependent on ST2 expression in Tregs and IL-33 treatment reprogrammed Tregs toward a more cytotoxic and inflammatory phenotype rather than simply enhancing their immunosuppressive activity. In contrast, the authors found that endogenous IL-33/ST2 signaling in Tregs promoted tumor growth. Thus, increased IL-33 availability may not necessarily enhance Treg-mediated immunosuppression and could instead alter Treg function toward a more inflammatory phenotype under certain conditions. Furthermore, the outcome of IL-33/ST2 signaling may depend not only on the amount of IL-33 available or on the type of cell that expresses ST2, but also on the functional state induced by IL-33/ST2 signaling within each responding population. Further studies are therefore needed to determine whether the increased extracellular IL-33 associated with the *Ala* IL-33 isoform alters the phenotype and suppressive functions of tumor-infiltrating ST2+ Tregs.

An additional consideration when interpreting the effects of tumor-derived IL-33 is the substantial discrepancy between our findings and those reported by *Zhang et al.*^29^. While we observed that tumor cell-derived IL-33 suppressed tumor growth, they reported that IL-33 overexpression increased MC38 proliferation *in vitro* and promoted tumor growth *in vivo*. Their *in vitro* assessment of cell proliferation was, similar to our study, based on cell counting and WST-1 cell proliferation assay, making differences in the general approach to assessing proliferation an unlikely explanation for the divergent findings. Several methodological differences between the two studies may nevertheless be relevant. First, whereas *Zhang et al.* employed a pcDNA3.1-based expression system to overexpress full-length IL-33, we used lentiviral vectors to introduce our IL-33 isoforms into MC38 cells, resulting in higher transgene expression. The higher levels of IL-33 expression in our model may result in differences in the local availability of IL-33 and consequently alter its effects on the tumor microenvironment. Second, we injected 0.5×10^6^ cells per flank, whereas *Zhang et al.* injected 2×10^6^ cells per flank. Although increased numbers of parental MC38 cells have previously been shown to accelerate subcutaneous tumor growth in WT mice^49^, whether this relationship also applies to IL-33-overexpressing MC38 cells is unclear. In our hands, injection of increasing numbers of MC38-*Il33* cells resulted in further reduction of tumor growth (data not shown). This suggests that differences in cell numbers alone are unlikely to explain the contrasting phenotypes observed in the two studies. Third, *Zhang et al.* did not provide information on tumor growth kinetics over the course of the experiment, limiting direct comparisons of tumor growth dynamics between the two studies. Fourth, *Zhang et al.* detected IL-33 in both parental and vector-control MC38 cells using multiple approaches, including qPCR and western blotting. Although the authors appeared to use the IL-33 antibody from the same commercial supplier as in our study, they did not report the antibody clone. This distinction may be relevant, since several commercially available IL-33 antibodies have been reported to exhibit nonspecific binding^23^. Indeed, our cytospin-based immunohistochemical analysis revealed a small number of parental and *Mock* MC38 cells showing positive IL-33 staining, whereas endogenous IL-33 was not detected by qPCR or western blotting, suggesting that this staining may represent nonspecific antibody binding. Consequently, differences in the endogenous IL-33 status of the parental cell lines, together with differences in cell origin and in the used IL-33 expression system, could substantially alter the baseline IL-33/ST2 signaling environment and thereby contribute to the contrasting phenotypes observed in the two studies. Taken together, our findings support a model in which the biological effects of tumor-derived IL-33 are determined by the balance between intracellular retention and extracellular availability of IL-33, as well as the composition of ST2-expressing immune populations. Increased extracellular availability of IL-33 was associated with enhanced frequencies of IFN-ɣ-producing cells and improved tumor control, suggesting that tumor-derived IL-33 can promote a type 1 immune response under these conditions. At the same time, the concurrent presence of ST2-expressing Tregs indicates that increased IL-33 availability may also promote their activation.

These results are particularly relevant for therapeutic strategies targeting the IL-33/ST2 pathway. Tumor-derived IL-33 has been shown to enhance responses to immune checkpoint blockade in MC38 tumors^34,39^, providing a rationale for therapeutic approaches aimed at increasing IL-33-mediated anti-tumor immunity. In contrast, other studies reported improved anti-PD-1 responses following inhibition of the IL-33/ST2 signaling pathway^44^, indicating that blockade of the same inflammatory pathway can be beneficial under different conditions. These apparently opposing therapeutic effects are consistent with the distinct cellular effects of ST2 signaling described above and suggest that systemic activation or inhibition of IL-33/ST2 signaling may not adequately account for the spatial and cellular complexity of the pathway. Our findings raise the possibility that selectively enhancing the local extracellular availability of IL-33 within the tumor may still represent a possible approach to boost anti-tumor immunity.

## Material and Methods

### Mice

All animal experiments were performed at the University of Bern, have been approved by the Cantonal Veterinary Office of Bern and are in accordance with Swiss Federal regulations. C57BL/6J mice were used as WT mice. *Il1rl1^tm1Anjm^*(*Il1rl1*^-/-^ or *St2*^-/-^) were generated by A.N. McKenzie^50^. In all experiments, non-randomized groups of 7-week-old mice were used and kept under SPF conditions.

### Cell culture and *in vitro* cell line characterization

MC38 (derived from C57BL/6 murine colon adenocarcinoma cells^51^) cell lines have previously been stably transfected with lentiviral vectors encoding three isoforms of IL-33 and a *Mock*. The MC38 cells were ordered from Kerafast (Boston, Cat#ENH204, Lot#011317). The adherent cells were cultured in growth medium (DMEM Sigma D6546, 10% FCS, 1mM non-essential amino acids, 2mM glutamine, 1mg/ml gentamycin, 0.01M HEPES, 1x Pen/Strep) in T75 flasks (Sarstedt, Cat#PW83.3911). To ensure selection of the transduced cells, the medium was supplemented with 8µg/ml puromycin (Sigma, Cat#P8833). Upon reaching 80% confluency, the cells were split by washing with PBS and detaching with trypsin (1x Trypsin-EDTA) for 3 min at 37°C. The cells were then transferred into new T75 flasks for maintenance or T175 (Sarstedt, Cat#PW833912) for further growth.

### Cytospin

Cultured cells were washed with PBS and detached from the surface using 1x Trypsin-EDTA, followed by an additional washing step. Cells were counted and resuspended in PBS with 10% FBS to a concentration of 1×10^6^ cells/ml. Slides were pre-coated with 50 µl PBS-10% FBS and centrifuged at 1500 rpm for 4 min at high acceleration. 50 μL of the prepared cell suspension were loaded into CytoSpin units and centrifuged at 400 rpm for 4 min at low acceleration. Slides were fixed in 4% Paraformaldehyde for 5 min and air-dried at room temperature. Prepared slides were sent for staining with murine IL-33 antibody (1:600 dilution).

### Detection of integrated IL-33 by PCR

Genomic DNA (gDNA) was extracted from 1×10⁶ MC38 cells that were 80% confluent. Cells were trypsinized and resuspended in 200 μL tissue lysis buffer, followed by the addition of Proteinase K (Roche, Cat# 3115801001) at 100 μg/ml. The mixture was incubated in a Thermo-Shaker at 600 rpm and 55°C for 30 min, followed by an inactivation step at 95°C for 20 min at 600 rpm. Each PCR reaction was prepared using a total volume of 20 μL, containing 5 μL of 5x buffer, 1 μL of dNTPs, 1 μL each of forward and reverse primers, 1.5 μL of MgCl₂, 0.25 μL of Taq polymerase, and 2 μL of gDNA. PCR amplification was performed under the following conditions: an initial denaturation at 95°C for 5 min, followed by 35 cycles of denaturation at 95°C for 30 sec, annealing at 55°C for 30 sec, and extension at 72°C for 2 min, with a final extension step at 72°C for 5 min. A total of 10 μL of the PCR product was combined with 3 μL of loading dye and loaded onto a 1% agarose gel. A 1 kb DNA ladder (cat# G7541) was used as a molecular weight marker. Gel electrophoresis was run at 130V for 45-60 min.

### Sanger sequencing

Sequencing of *FL* and *Ala* cells was performed by centrifugation using the Wizard SV Gel and PCR Clean-Up System (Promega, cat# A9281), following the manufacturer’s instructions. DNA concentration was measured using NanoDrop. The samples were diluted to a final concentration of 4 μM using nuclease-free water (cat# P1193). The purified DNA samples were premixed with the corresponding sequencing primers and sent to Microsynth for sequencing. The sequencing results were analyzed using SnapGene Viewer (version 5.3.2).

### Cell fractionation

Cells were seeded and harvested at 80% confluency. Cells were washed with PBS, trypsinized and collected in a 1.5 ml Eppendorf tube. After a brief centrifugation at 16,000g for 10 sec, the supernatant (SN) was discarded and the pellet was resuspended and triturated in 900 μl of 0.1% NP-40 in PBS buffer. A 300 μl aliquot of the lysate was set aside as the “whole cell lysate” fraction, and 100 μl of 4x Lämmli sample buffer (cat# 1610747) supplemented with 1:10 2-Mercaptoethanol (cat# 8.0574) was added before keeping it on ice. The remaining 600 μl of lysate was centrifuged at 16,000g for 10 seconds, and a 300 μl aliquot of the supernatant was collected as the “cytosolic fraction”, to which 100 μl of 4x Lämmli buffer supplemented with 2-Mercaptoethanol was added before storing on ice. The remaining supernatant was discarded, and the pellet was resuspended in 1 ml of ice-cold 0.1% NP-40 in PBS, followed by another centrifugation at 16,000g for 10 sec. The supernatant was discarded and the pellet was resuspended in 180 μl of 1x Lämmli buffer supplemented with 2-Mercaptoethanol, designated as the “nuclear fraction”. The nuclear fraction and whole cell lysate were sonicated using an ultrasonic homogenizer (Bandelin, Sonopuls GM3200) at 10% amplitude, with two pulses of 5 sec each. All samples (whole cell lysate, cytosolic fraction, and nuclear fraction) were boiled at 95°C for 5 min and stored at 4°C.

### Western blot

The TGX Stain-Free FastCast Polyacrylamide 12% Kit (Bio-Rad, cat# 1610185) was used to prepare gels and the Kaleidoscope Standard (Bio-Rad, cat# 1610375) was used as a molecular weight marker. The gels were running at 120V for 60–90 min, followed by UV activation for 1 min using the ChemiDoc imaging system (Bio-Rad). For protein transfer, the PVDF membrane was incubated in 100% methanol, washed in TBS and assembled into a transfer stack. Protein transfer was performed using the Trans-Blot Turbo Transfer System (Bio-Rad) with the High MW program (1.3A, 25V) for 10 min. After, the membrane was blocked with 5% non-fat dry milk (NFDM) in TBS-T (Tris-buffered saline with Tween-20) for 60 min at room temperature. The membrane was then incubated overnight at 4°C with the polyclonal primary antibody (goat anti-mouse IL-33, RCD Systems, cat# AF3626) diluted 1:500 in 5 ml of 5% NFDM in TBS-T. The next day, the membrane was washed three times for 5 min with 10 ml TBS-T, followed by incubation with the secondary antibody (Goat IgG HRP-conjugated, RCD Systems, cat# HAF109), diluted 1:1000 in 5 ml of 5% NFDM in TBS-T for 2 h at room temperature. The membrane was then washed as described above and given a final 10 min wash with 10 mL TBS. To verify the purity of the cytoplasmic fraction, the membrane was probed for the cytoplasmic protein GAPDH using the primary antibody Anti-GAPDH, clone 6C5 (Merck Millipore, cat# MAB374), diluted 1:5000 in 5 ml NFDM in TBS-T. For the nuclear fraction, a separate membrane was probed for the nuclear protein NUP85 using the monoclonal anti-NUP85 antibody (RCD Systems, cat# MAB5976), diluted 1:2000 in 5 ml NFDM in TBS-T. Detection was performed by washing the membranes as described above, incubating them with ECL reagent (1:1 ratio) for 5 min and imaging on the ChemiDoc MP system using manual acquisition or optimal auto-exposure mode.

### Apoptosis induction

Cells were seeded in a 24-well plate in 500 μl of complete DMEM (high glucose). After 48 h, a concentration of 2.25 mM DL-Dithiothreitol (DTT) solution (Sigma, cat# 43816-10mL) was added to the medium to determine apoptosis induction. Supernatants were collected in Eppendorf tubes immediately before DTT administration and 24 h post-treatment. The collected supernatants were centrifuged at 16,000g for 10 s, transferred into fresh 1.5 ml Eppendorf tubes, and stored at -80°C for further analysis. In parallel, cells were retained for FACS analysis to assess apoptosis induction. For this, the cell pellet was resuspended in 100 μl of Annexin binding buffer (Biolegend cat# 422201) and transferred to a 96-well plate for staining. Cells were centrifuged at 300g for 5 min, the supernatant was discarded, and the pellet was washed twice with 100 μl of Annexin binding buffer. Cells were then incubated with the antibody mix, consisting of Annexin V (Biolegend cat# 640918) (1:100) and 7AAD (Biolegend cat#420404) (1:100) diluted in Annexin binding buffer, for 15 min at room temperature in the dark. Staining with Annexin V and 7-aminoactinomycin D (7-AAD) enables differentiation between viable cells (Annexin V-negative, 7-AAD-negative), early apoptotic cells (Annexin V-positive, 7-AAD-negative), and late apoptotic or necrotic cells (Annexin V-positive, 7-AAD-positive).

### Assessing IL-33 Release After Necrosis Induction

Necrosis was induced through multiple freeze-thaw cycles. Cells were trypsinized using Trypsin-EDTA, centrifuged at 300g for 5 min, and the supernatant was discarded. The resulting pellet was resuspended in 1 ml of PBS. For the freezing step, the microcentrifuge tube containing the resuspended cells was submerged in liquid nitrogen for 2 min until fully frozen. The thawing step was then performed by immediately transferring the frozen tube to a 37-42°C water bath and allowing the cells to thaw completely for 1 min. Once thawed, the tube was vortexed to ensure cell disruption and homogenization. The freeze-thaw process was repeated for a total of three cycles to maximize necrosis induction. After the final thawing step, the tube was centrifuged at 300g for 5 min. Cells were immediately analyzed by flow cytometry to assess necrosis induction, while the supernatants were collected and used in ELISA to detect IL-33 secretion.

### Enzyme-linked Immunosorbent Assay (ELISA)

ELISA was performed using the Mouse IL-33 ELISA Kit from RCD Systems (cat# DY3626-05, lot# P141478), following the manufacturer’s protocol. Standard ELISA plates (Nuclon, Thermo Scientific, cat# 442404) were coated with a goat anti-mouse IL-33 capture antibody and incubated overnight at 4°C. Undiluted samples were added to the plates in duplicate, followed by incubation with a biotinylated goat anti-mouse IL-33 detection antibody and streptavidin conjugated to horseradish peroxidase (HRP). The optical density (OD) was measured using the Tecan Infinite 200 PRO at 450 nm.

### *In vivo* cell growth and subsequent analysis of immune cells

To prepare the MC38 cell lines for subcutaneous injections, the cells were harvested when they reached a confluency of 80%. For this, the cells were washed with PBS, trypsinized to detach the adherent cells, centrifuged and washed with PBS. The cells were then counted manually with a Neubauer chamber and resuspended in PBS at a concentration of 2.5×10^6^ cells/ml (corresponding to 0.5 ×10^6^ cells/200µl per injection). Mice were injected subcutaneously into both flanks, and tumor growth was measured 2-3x per week using a caliper. Tumor volume was calculated by multiplying the length, width and height of each tumor. The mice were euthanized 18-19 days post injection and tumors and spleens were excised for subsequent analysis of the immune cells. Splenic immune cells were obtained by smashing the spleen through a 70 µm filter. Tumor immune cells were isolated with a digestion mixture consisting of RPMI 1640 media (Sigma, R0883), 13 WU Liberase (Roche, Cat#05401020001) and 10mg/ml DNase I (Roche, Cat#10104159001) for 40min at 37°C, followed by filtration trough a 70µm filter. After a washing step, spleen and tumor cell suspensions were lysed (red blood cell lysis) with ACK lysis buffer (5min), followed by resuspension in PBS for subsequent flow cytometry analysis.

### Flow cytometry

Isolated cells were stained using the marker-specific fluorochrome-labeled antibodies indicated in Supplementary Table 1. For the fixation of samples containing only surface antibodies, a Cytofix/Cytoperm kit (BD Biosciences, Cat#554715) was used, while for samples containing intracellular proteins a Foxp3/Transcription staining buffer kit (ThermoFisher, Cat#00-5523-00) was used. Cells isolated from IL-33 reporter mice were not fixed, to retain the GFP-Il33 signal. To assess intracellular IFN-ɣ or TGF-β, cells were first stimulated with 1µg/ml ionomycin (Sigma, Cat#I0634), 10ng/ml phorbol-12-myristate 13-acetate (PMA, Sigma, Cat#P1585) and 1x Brefeldin A (Biolegend, Cat#420601). Samples were acquired using the Aurora 5L (Cytek) and analyzed using the FlowJo software (v10.10.1, Waters Biosciences).

## Statistics

Statistical tests were performed with GraphPad Prism (v11.0.1) for Windows. For single comparison of normally distributed data, unpaired t test with Welch’s correction was used. To compare the means of two groups or more, 1- or 2-way ANOVA with Tukey’s or Šidák’s multiple comparisons test was used. For all statistical analyses: *:P<0.05; **P<0.01; ***P<0.001; ****P<0.0001.

## Author contributions

AH, IE, IK, SE, VPV and MHW conducted experiments and acquired and analysed data. KS and CN conducted experiments. RG analysed and discussed data. AH, VPV, MHW and PK designed and supervised research studies. AH and PK wrote the manuscript. All authors approved the final version of the manuscript.

## Acknowledgements

We thank the Translational Research Unit of the Institute of Tissue Medicine and Pathology and the FACS facility of the University of Bern for excellent technical support and advice.

