## Supplementary material for "Extracellular tumor-derived IL-33 promotes ST2-dependent tumor control": Suppl. Figures

### Supplementary Information

#### Supplementary Figure 1

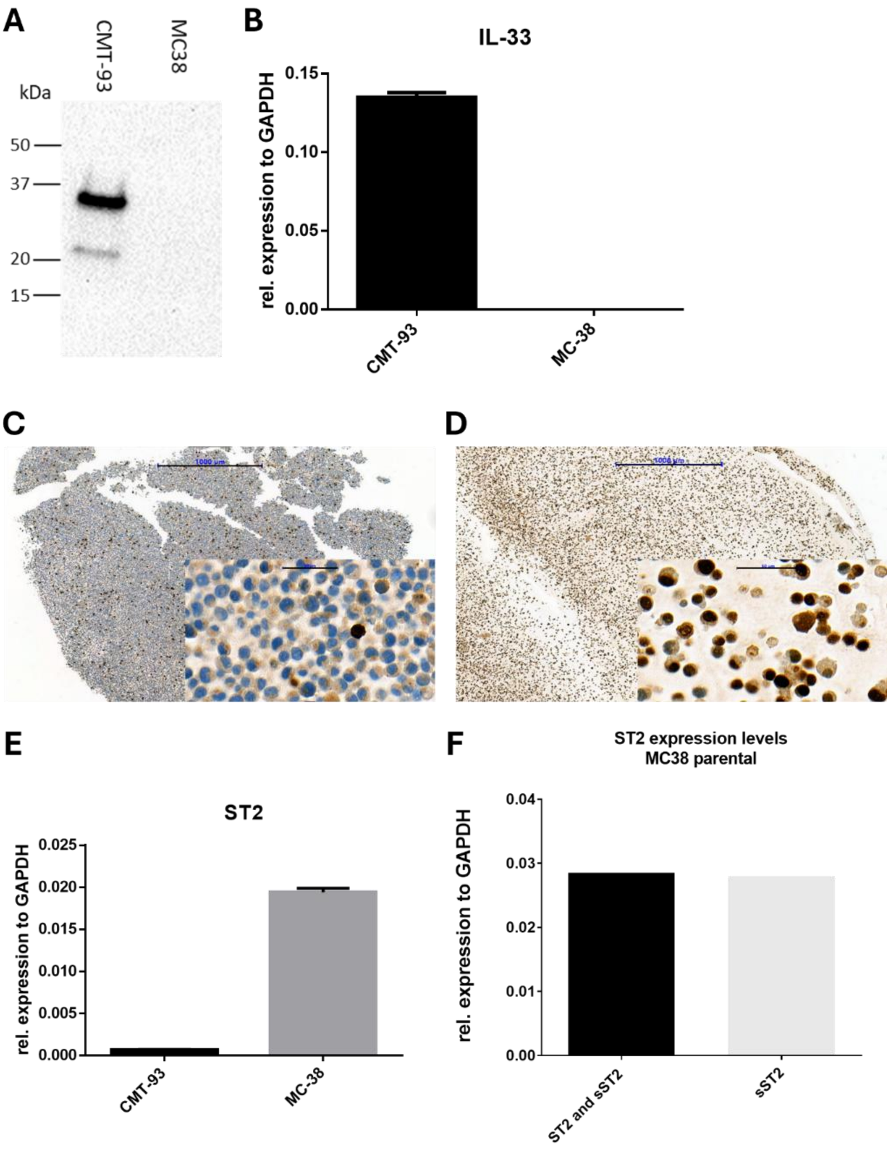


***Supplementary Figure 1: Expression of IL-33 and ST2 in the parental MC38 and CMT-93 cell line.*** *(A) Western blot analysis of IL-33 in CMT-93 and MC38 cells. IL-33 expression was below the limit of detection in MC38 cells. (B) qPCR of Il33 in CMT-93 and MC38 cells. Cell culture pellet of (C) MC38 and (D) CMT-93 cells stained for IL-33 (brown). Scale bar overview: 1000µm, scale bar inlay: 50µm. (E) qPCR of St2 in CMT-98 and MC38 cells. The primer set used detects the soluble (sSt2) and receptor form of St2. (F) qPCR of St2 on MC38 cells. Left: primers detect St2 and sSt2, right: primers detect only sSt2. (A) Blot representative of 3 independent experiments. (E-F) Blot representative of 2 technical replicates.*

**Supplementary Figure 2**

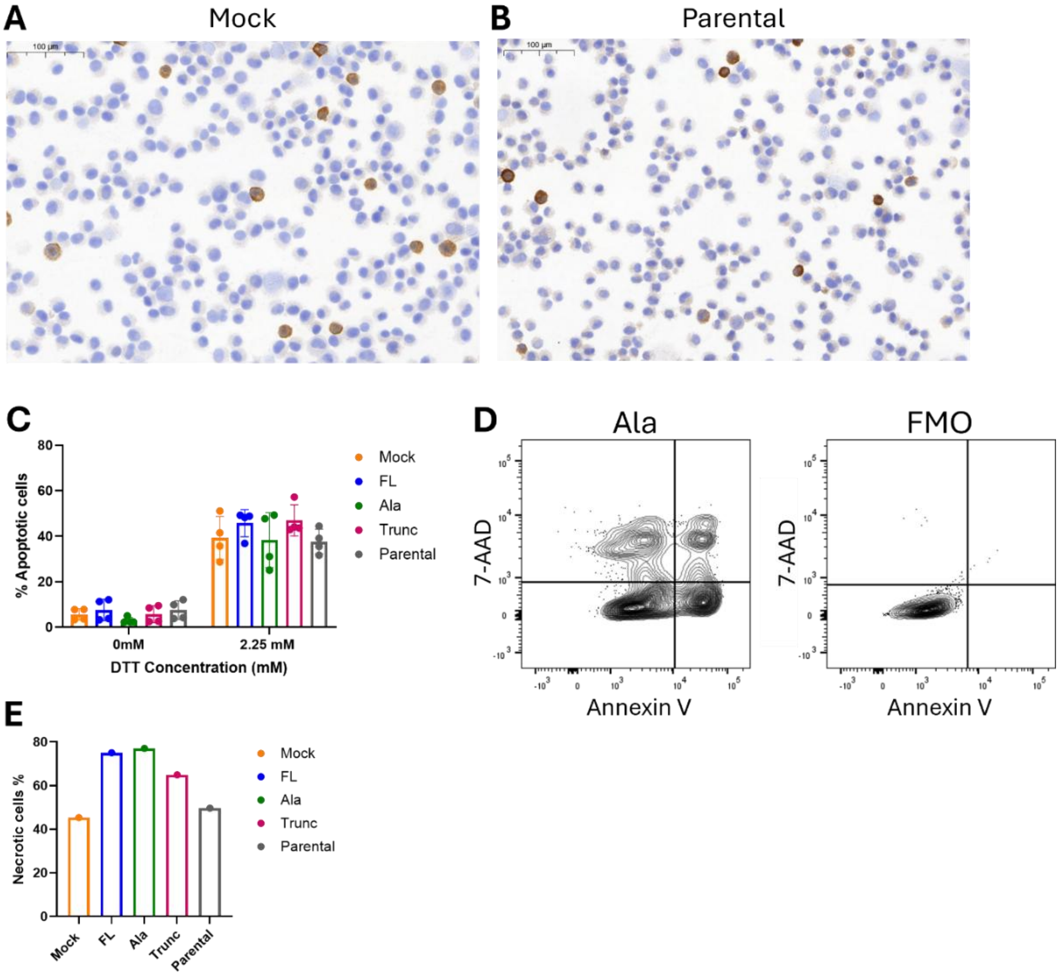


***Supplementary Figure 2: In vitro characterization of MC38-Il33 cells.*** *Zoom-out of the (A) Mock and (B) Parental MC38 cell lines prepared using the cytospin method and stained for IL-33 (brown). IL-33+ cells account for 3-5% for both cell lines. (C) A concentration of 2.25 mM DTT induced comparable levels of apoptosis in 40-50% of cells across the MC38-Il33 cell lines. The percentage of apoptotic cells of MC38-Il33 cells before (0h) and after (24h) DTT-induced apoptosis was analyzed using flow cytometry analysis. (D) Gating strategy to identify apoptotic cells (7-AAD negative, Annexin V positive). Left: representative plot from Ala cells, right: FMO as a negative control. (E) Necrosis was induced via freeze-thaw cycles in the indicated MC38 cells. Percentage of necrotic cells (7-AAD positive, Annexin V positive).*

#### Supplementary Figure 3

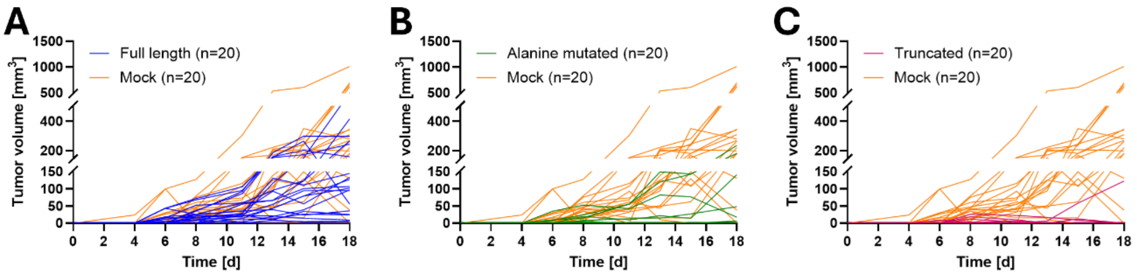


***Supplementary Figure 3: High release of IL-33 decreases MC38 tumor growth.*** *MC38-Il33 cell lines were injected subcutaneously into both flanks of WT mice. Tumor volume was measured every 2-3 days with a caliper. Individual tumor growth of (A) Full length and Mock, (B) Alanine mutated and Mock and (C) Truncated and Mock. Data pooled from 2 independent experiments, N=10 mice per group.*

**Supplementary Figure 4**

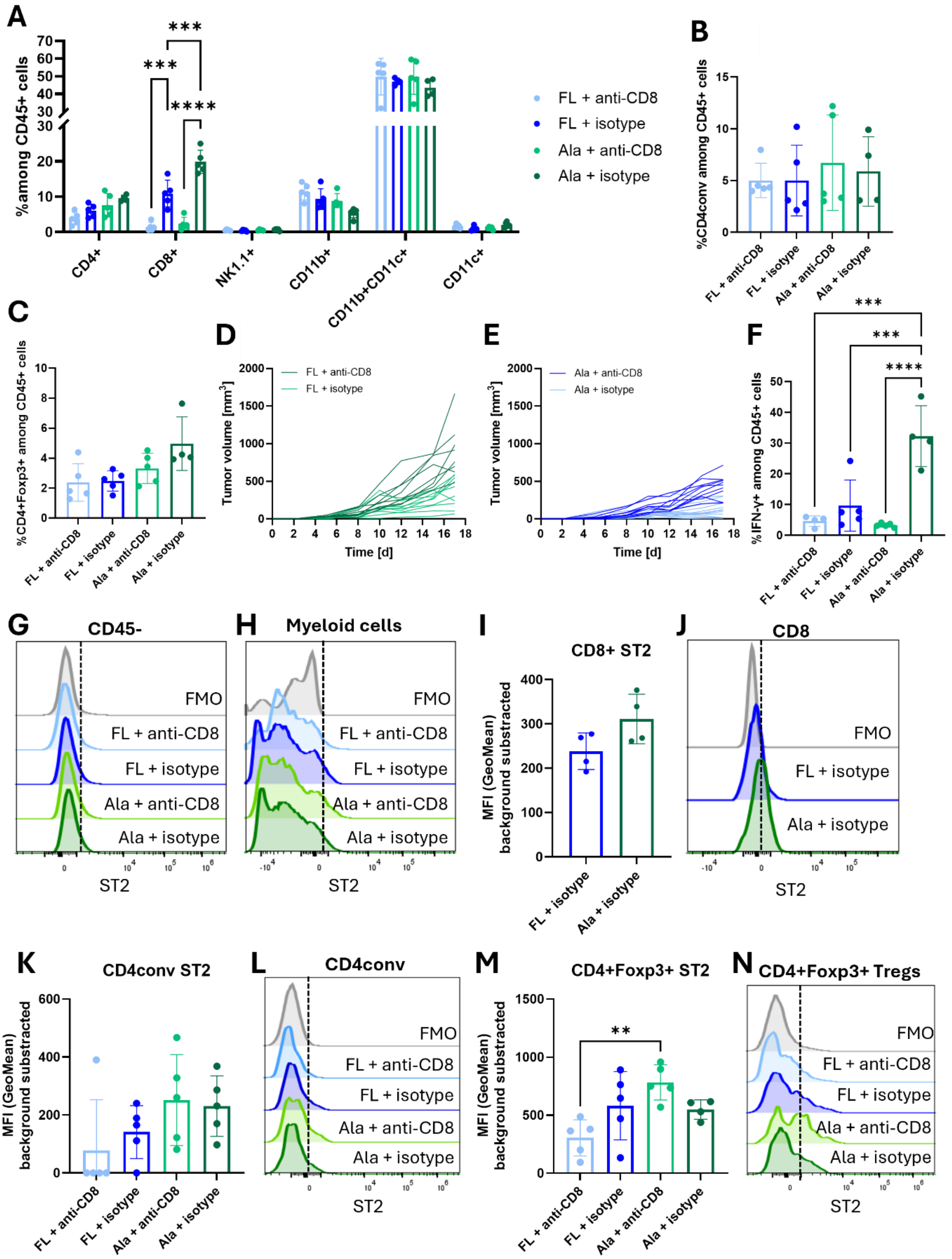


*__Supplementary Figure 4: IL-33-dependent tumor growth is in part controlled by CD8+ T cells.__ WT mice were subcutaneously injected into both flanks with FL or Ala MC38 cells in combination with either depleting anti-CD8 antibody or isotype control. At the endpoint, tumors were excised and digested for analysis of tumor-infiltrating immune cells. (A) Frequencies of indicated tumor-infiltrating immune cell subsets among CD45+ immune cells. Frequencies of (B) conventional CD4+ T cells (CD4conv) and (C) CD4+Foxp3+ Tregs among CD45+ cells. Individual tumor growth of (D) FL MC38 cells and (E) Ala MC38 cells treated with depleting anti-CD8 antibody or isotype control. (F) IFN-ɣ+ among CD45+ cells. Representative graphs of MFI of ST2 expression of (G) CD45- and (H) myeloid cells (CD11c+, CD11b+ and CD11b+CD11c+). The dotted line represents the cut-off for ST2 expression. (I) MFI with subtracted background of ST2*

*expression on CD8+ T cells and (J) representative MFI graphs. (K) MFI with subtracted background of ST2 expression on CD4conv and (L) representative MFI graphs. (M) MFI with subtracted background of ST2 expression on CD4+Foxp3+ Tregs and (N) representative MFI graphs. (A-N) Data from 1 experiment, N=4-5 mice per group (A-C, F, I, K, M) 1-way ANOVA with Tukey's multiple comparisons test, (I) Unpaired t test with Welch's correction (ns). *: P < 0.05; **: P < 0.01; ***: P < 0.001; and ****: P < 0.0001.*

#### Supplementary Figure 5

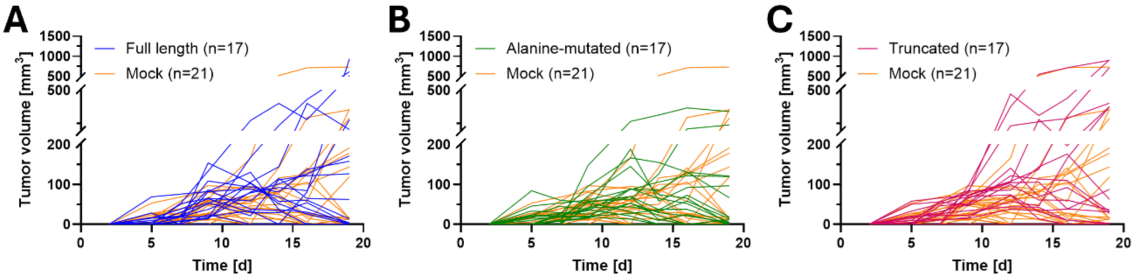


***Supplementary Figure 5: Ablation of St2 abrogates the anti-tumor effect of IL-33.*** *MC38-Il33 cell lines were subcutaneously injected into both flanks of $St2^{-/-}$ mice. Tumor volume was measured every 2-3 days with a caliper. Individual tumor growth of (A) Full length and Mock, (B) Alanine mutated and Mock and (C) Truncated and Mock. Data pooled from 2 independent experiments, N=10-12 mice per group.*

**Supplementary Table 1**

***Supplementary Table 1: Antibodies used for flow cytometry, including their conjugate, clone name, catalog number and source.***

| Specificity | Conjugated to | Clone Name | Catalog number | Source |
|---|---|---|---|---|
| CD45.2 | AF700 | 104 | 109821 | Biolegend |
| Viability dye | Fixable blue | | L34962 | Thermofisher |
| CD45.2 | BV570 | 30-F11 | 103136 | Biolegend |
| CD4 | Pacific Blue | RM4-4 | 116008 | Biolegend |
| CD8a | BV711 | 53-6.7 | 100747 | Biolegend |
| CD11c | RB705 | N418 | 570627 | BD (Waters Biosciences) |
| CD11b | RY703 | M1/70 | 571473 | BD (Waters Biosciences) |
| NK-1.1 | BUV805 | PK136 | 741926 | BD (Waters Biosciences) |
| GR-1 | PE-Cy7 | RB6-8C5 | 108416 | Biolegend |
| B220 | Pe/Cy5 | RA3-6B2 | 103210 | Biolegend |
| ST2 | APC | DIH9 | 145306 | Biolegend |
| CD4 | APC-Cy7 | RM4-5 | 100525 | Biolegend |
| IFN-ɣ | Spark NIR™ 685 | XMG1.2 | 505862 | Biolegend |
| FoxP3 | FITC | FJK-16s | 11-5773-82 | eBioscience |
| Annexin V | Pacific Blue | | 640918 | Biolegend |
| 7AAD | | | 420404 | Biolegend |
